# Role of Age in Mediating the Association Between the Vaginal Microbiota and Preterm Birth

**DOI:** 10.1101/2025.01.27.635059

**Authors:** Yun Xie, Qi Wang, Dan Li, Fengan Jia, Fan Chang, Zhen Zhang, Yanmei Sun, Shiwei Wang

## Abstract

The vaginal microbiota plays a crucial role in vaginal health and pregnancy outcomes. However, the influence of maternal characteristics, especially age, on the relationship between the vaginal microbiota and preterm birth is poorly understood. This study quantifies the effects of race, maternal age, and gestational age on vaginal microbiota composition. Our results demonstrate stronger associations between preterm birth and a less optimal vaginal microbiota in White and Asian women compared to Black or African American women. The microbiota difference between preterm and term births was more pronounced in older reproductive-age women, with *Lactobacillus* species increasing with age only in term births. Additionally, lower alpha diversity was observed in term pregnancies compared to preterm during both early and late pregnancy, particularly in women aged above 25 years. These findings highlight the mediating role of age in the relationship between the vaginal microbiota and preterm birth.

## Introduction

The vaginal microbiota (VMB) is a critical component of the female reproductive system, playing an essential role in maintaining vaginal health and influencing pregnancy outcomes^1^. The dominance of *Lactobacillus* species in the VMB is a key indicator of overall vaginal health. Protective *Lactobacillus* contribute to microbial balance by producing lactic acid, which helps maintain a low vaginal pH, suppresses the growth of pathogens, and supports immune modulation. In contrast, many non-*Lactobacillus* species, such as *Gardnerella vaginalis, Sneathia amnii*, and *Lachnospiraceae* BVAB1, produce virulence factors and are linked to various diseases in the female lower reproductive tract.

Preterm birth (PTB), defined as delivery before 37 weeks of gestation, is a major global health concern, accounting for approximately 10% of all live births and contributing to high rates of neonatal morbidity and mortality^2^. The causes of PTB are multifactorial, involving genetic, environmental, and microbial factors^3^. Among these, the role of the VMB has received significant attention. Studies have shown that certain bacterial taxa are associated with increased risk of PTB, including *G. vaginalis, S. amnii*, and other dysbiosis-related species^1^. Conversely, a VMB dominated by health-associated *Lactobacillus* species, e.g., *L. crispatus, L. jensenii*, and *L. gasseri*, is generally linked to better pregnancy outcomes. *L. iners* is the only *Lactobacillus* species whose role in overall vaginal health remains unclear^4^. Recent studies suggest that a vaginal microbiome dominated by *L. iners* may represent an intermediate or transitional state of microbial composition^5^.

The composition of the VMB is influenced by several factors, including race and gestational stage^1^. Studies have consistently reported that Black/African American (B/AA) women are more likely to have VMBs characterized by higher microbial diversity and increased prevalence of dysbiosis-associated taxa compared to White women^6^. Gestational age also plays a critical role, as during pregnancy, the VMB undergoes substantial compositional and functional changes, marked by an increased dominance of *Lactobacillus* species and a reduction in taxa associated with dysbiosis^7^. This change is thought to represent a protective mechanism that supports healthier delivery outcomes.

Despite advances in VMB research, the interaction between the VMB and maternal characteristics in pregnancy outcomes remains unclear. Most studies on VMB and PTB are cross-sectional and often neglect confounding factors such as race and gestational age^8^. Additionally, studies on the association between maternal age and the VMB primarily compare the VMB in postmenopausal and reproductive-age women^9^, leaving the role of maternal age in the association between the VMB of reproductive-age women and PTB poorly understood.

In this study, we sought to address these gaps by analyzing a large, high-quality dataset of VMB samples collected from pregnant women across diverse racial groups, ages, and gestational ages. Our data elucidate the influence of race, age, and gestational age on the association between the VMB and PTB by integrating data from multiple cohorts and employing robust statistical approaches. Our findings underscore more pronounced differences in the VMB between PTB and term births in older women of reproductive age. The data also highlight that, in term births, the relative abundances of *L. crispatus* and *L. jensenii* increase with age, while the abundance of dysbiosis-associated taxa decreases, but similar trends are not in PTBs.

## Methods

### Inclusion and exclusion criteria

Publications were searched in the PubMed database using a strategy that incorporated three key terms: (“vaginal microbiota” OR “vaginal microbiome”) AND (16S rRNA) AND (preterm OR premature). The inclusion criteria were as follows: 1) Raw 16S rRNA sequencing data were publicly available for download from the Sequence Read Archive (SRA); 2) The metadata of participants, i.e., race, age, gestational age, and pregnancy outcome, were clearly specified; 3) The study was an original research article. The exclusion criteria included: 1) Studies not related to the VMB; 2) Studies with fewer than 30 samples; 3) Participants under the age of 16 were excluded.

### Data preprocessing

Following quality control, paired sequence reads were trimmed, merged, and human-origin reads were removed as outlined previously^10^, The resulting high-quality 16S rRNA amplicon sequences were then aligned to a refined 16S rRNA database described in the earlier research^10^. Taxa were included in the feature table only if their relative abundance reached at least 0.1% (or 0.01%) in 5% (or 15%) of the samples, as previously reported^10^. Samples with fewer than 5,000 total reads were excluded from further analysis.

### Removal of batch effect

Batch effects across cohorts were corrected using the sva package in R. Specifically, the feature table of 16S rRNA profiles was normalized using the CLR normalization and the ComBat function was applied to address batch effects.

### dbRDA analysis

The influence of multiple variables, i.e., race, age, gestational age, and pregnancy outcome, on the composition of the VMB was assessed using the dbRDA test implemented through the capscale function in R, with the parameter distance set to ‘bray’^11^. Statistical significance was evaluated using the marginal Adonis test, with the parameter ‘by’ set to “margin.” To ensure comparable degrees of freedom among these variables, age was categorized into three levels: ‘low,’ ‘medium,’ and ‘high,’ with cases evenly distributed across these groups. Similarly, gestational age was recoded into three categories.

### Alpha and beta diversity

The feature table derived from 16S rRNA profiles was normalized through rarefaction, matching the sequencing depth of the sample with the fewest reads (>5,000). To quantify alpha diversity, the Shannon index was computed using the R package vegan^11^, and group differences were analyzed via the two-sided Wilcoxon test. Beta diversity was visualized using the prcomp function in R to calculate the PCoA values. Group-level beta diversity differences were examined through PERMANOVA analysis using the ‘adonis2’ function provided in the vegan package.

### Differential abundance analysis

Differential abundance analysis was performed using the ALDEx2 package in R^12^. The adjusted P-values for differences in relative abundance were computed with the aldex.ttest function, employing the Mann-Whitney U test and subsequent Benjamini-Hochberg correction. Changes in relative abundance were quantified using the aldex.effect function, which calculated the median difference per feature between two conditions.

### Interaction between age and preterm birth in relation to bacterial taxa

A mixed-effects model was implemented using the lm function from the lme4 package, with the formula specified as Bacterium ∼ Age * Outcome. The bacterial abundances were the CLR normalized values of bacterial reads. Two sets of P-values were generated: one for the correlation between age and bacterial abundance, and another for the interaction effect between age and pregnant outcomes on bacterial abundance. Both sets of P-values were adjusted using the Benjamini-Hochberg correction to control for multiple comparisons.

### Institutional review board statement

Study protocols were approved by the Northwest Women’s and Children’s Hospital Institutional Review Board (IRB) under protocol number IRB 2024-100. Since all the publicly available datasets were obtained from the NCBI database (see details in Data availability), information regarding informed consent to participate can be found in the original publications associated with these datasets^13–16^.

## Results

### Overall profiles of the vaginal microbiota

This study analyzed 2,408 samples (Fig. S1) from four BioProjects (Fig. S2a)^13–16^, selected based on criteria described in the Methods. Alpha rarefaction analysis determined an appropriate threshold for total reads, showing no significant difference in observed taxa between rarefaction depths of 1,600 and 1,800 reads (Fig. S1). To ensure high-quality data, samples with fewer than 5,000 reads were excluded as previously described^10^, leaving 2,266 samples for analysis. Low-abundance and rarely observed taxa were also removed, resulting in 143 species-level taxa for further study.

The final dataset included samples mostly from the United States (Fig. S2b). These included 516 PTB and 1,750 term birth participants (Fig. S2c). The majority of participants were B/AA, followed by White and smaller proportions of Asian and other racial groups (Fig. S2d). The average gestational age was 160.2 ± 60.8 days, and the mean participant age was 28.2 ± 5.3 years (Fig. S2e, S2f). Batch effect correction was applied to ensure data comparability across cohorts (Fig. S3).

Using all samples without case matching, alpha diversity (Shannon index) was significantly lower in term birth VMBs (Fig. S4a). Beta diversity, visualized with PCoA and quantified using the Adonis test, indicated a significant association between VMB composition and PTB (Fig. S4b). Differential abundance analysis identified five *Lactobacillus* species, e.g., *L. crispatus* and *L. jensenii*, as more abundant in term birth VMBs (Fig. S4c). Dysbiosis-associated taxa, e.g., *S. amnii, Lachnospiraceae* BVAB1, and *G. vaginalis*, were more abundant in preterm VMBs. These findings align with prior studies^1^.

### dbRDA analysis on factors associated with preterm

The associations between the VMB and factors, i.e., race, PTB, and gestational age, have been explored^1^, but their relative impacts on VMB composition have been quantified by limited studies^17^. Additionally, the relationship between age and the VMB in reproductive-aged individuals has not been well defined. To address these gaps, distance-based redundancy analysis (dbRDA) was applied to visualize the effects of these factors on VMB composition and an Adonis test with a marginal model was used to quantify their impact. For comparability, age (Fig. 1a) and gestational age (Fig. 1b) were divided into three evenly distributed categories.

**Fig. 1.**
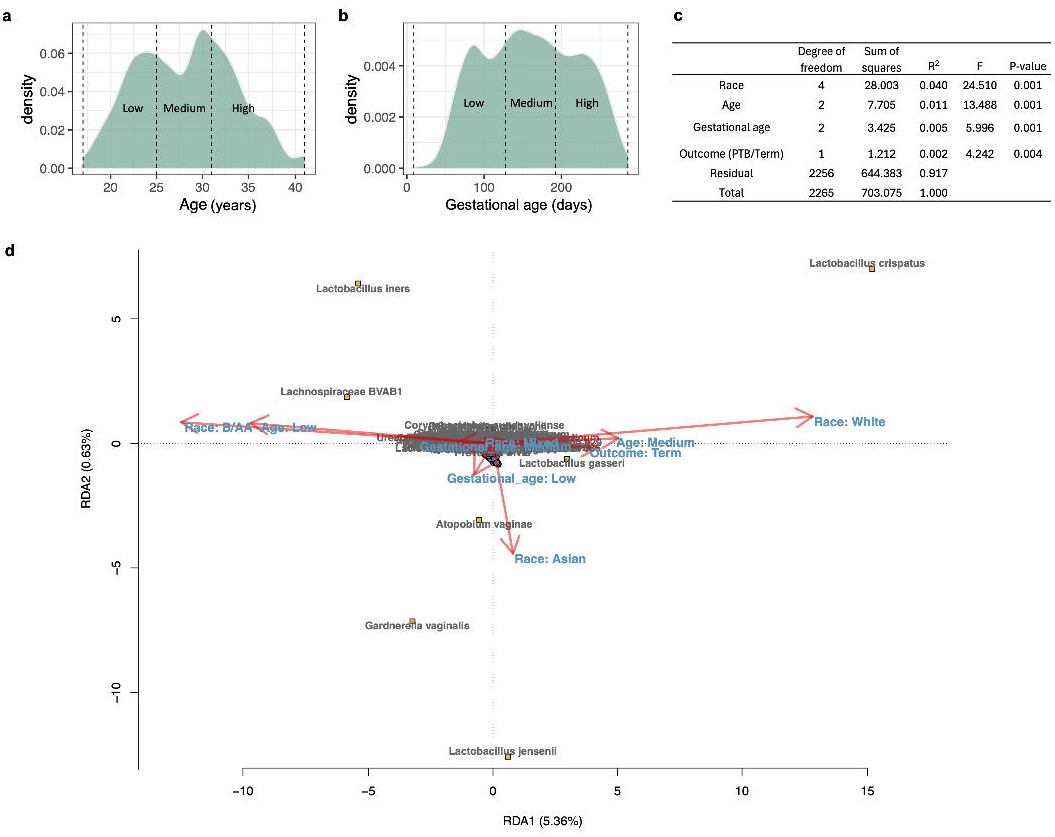
The composition of the vaginal microbiota (VMB) influenced by maternal factors. Age **(a)** and gestational age **(b)** divided into three categories, respectively, based on evenly distributed case numbers are shown. **(c)** Results of the Adonis test using a marginal model to quantify the effects of age, gestational age, and pregnancy outcome on VMB composition are illustrated. PTB represents preterm birth. **(d)** dbRDA analysis evaluating the influence of maternal factors on the composition of the VMB is shown.

Results of the Adonis test with a marginal model showed that race, age, gestational age, and birth outcome all significantly influenced VMB composition (Fig. 1c). Race accounted for the most variance, followed by age, gestational age, and birth outcome. VMBs from women aged 25.1-32 years, White women, or those with term births were enriched in *L. crispatus* and *L. gasseri* (Fig. 1d). Conversely, VMBs from women aged 17-25 years or of B/AA race were enriched in *L. iners* and *Lachnospiraceae* BVAB1. Samples collected earlier in pregnancy (arrow labeled ‘Gestational_age: Low’ in Fig. 1d) showed higher abundances of *A. vaginae* and *G. vaginalis*. These findings align with previous studies showing that White women have VMBs enriched with health-associated *Lactobacillus* species compared to B/AA women^6,15^, and that the VMB tends to enrich in *Lactobacillus* species and reduce dysbiosis-associated taxa during pregnancy^1,7^. Age was identified as the second most influential factor, with women aged 25.1-32 having more optimal VMB profiles.

### Association between the vaginal microbiota and preterm in different races

Some previous studies on the association between the VMB and PTB have been cross-sectional with limited control for confounding factors^8^. Although some studies matched samples based on race or ethnicity, age, and household income^18^, they lacked in-depth analyses of specific subgroups classified by race, maternal age, and gestational age. Here, VMBs from PTB and term birth cases were matched based on race, BioProject, and similar age and gestational age (Fig. S4d). The results showed that both alpha (Fig. S4e) and beta (Fig. S4f) diversities of the VMB remained significantly associated with PTB, but the significance level was obviously lower than in the unmatched analysis (Fig. S4a and S4b). Additionally, no taxa showed significant differences in relative abundance between PTB and term birth cases. These findings, together with the dbRDA analysis, suggest that race, age, and gestational age are confounding factors in studying the association between the VMB and PTB.

To exclude the impact of race, analyses were performed within specific racial groups. For each comparison, samples were matched by BioProject, age, and gestational age (Fig. S5a-c). The results showed that VMBs from White and Asian women, but not B/AA women, had significantly lower alpha diversity in term births compared to PTB (Fig. 2a). Consistent with previous research^17^, the beta diversity association with PTB was also stronger in White and Asian women than in B/AA women (Fig. 2b). In B/AA women, preterm birth was associated with higher levels of *Prevotella buccalis*, while term births in White women were enriched in *L. crispatus, L. jensenii*, and *L. kitasatonis* (Fig. 2c). In Asian women, term births showed higher levels of *L. crispatus, L. jensenii*, and *L. iners*, while preterm births had increased dysbiosis-related taxa, such as *G. vaginalis* and *Streptococcus anginosus*. However, most Asian participants were aged 32.1-42, which may have influenced these results, as discussed further below.

**Fig. 2.**
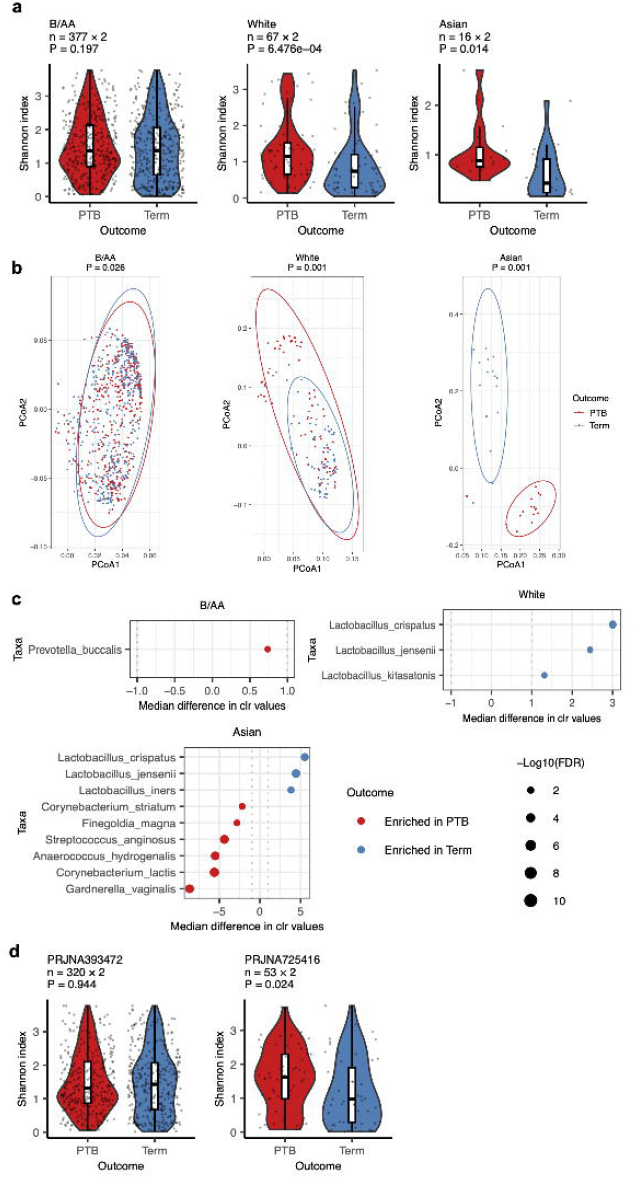
The impact of race on the association between the VMB and PTB. The VMB samples were categorized into three racial subgroups, i.e., Black or African American (B/AA), White, and Asian. In each subgroup, PTB and term birth samples were case-matched by BioProject, age, and gestational age (see Fig. S5). **(a)** Differences in alpha diversity between PTB and term birth within each racial group are shown. **(b)** Beta diversity differences visualized via principal coordinate analysis (PCoA) are illustrated. **(c)** Differential abundances of taxa in the VMB of PTB and term birth in the three racial groups are visualized by dot plots. **(d)** Differences in alpha diversity between PTB and term birth in B/AA women in two individual cohorts are depicted. Statistical analyses were conducted using a two-sided Mann-Whitney U test for alpha diversity and the Adonis test for beta diversity. Changes in relative abundance were tested using the ALDEx2 package in R, and quantified by the per-taxon median difference between conditions. FDRs were calculated with the Benjamini-Hochberg correction applied to the Mann–Whitney U test. Lines in the boxplots represent maximum, 75% quantile, median, 25 quantile, and minimum values from top to bottom.

Further analysis of B/AA women using case-matched VMBs (Fig. S5d and S5e) from two BioProjects, PRJNA393472 and PRJNA725416, showed lower alpha diversity in term birth women only in PRJNA725416 (Fig. 2d). However, the P-value for alpha diversity analysis in B/AA women of PRJNA725416 was less significant than in White women (Fig. 2a), and no taxa in B/AA women showed differential abundance linked to preterm birth in PRJNA725416. These results suggest inconsistent findings across cohorts for B/AA women, but PTB-associated differences were less pronounced in B/AA women compared to White and Asian women.

### Association between the vaginal microbiota and preterm mediated by age

The VMBs of different racial groups were further analyzed by stratifying women into age groups, with samples matched by BioProject, age, and gestational age (Fig. S6a for B/AA and S6b for White women). Interestingly, in B/AA women, alpha diversity associated with PTB was not significant in those aged 17-25 but became significant in the 25.1-32 age group and was most significant in those aged 32.1-42 (Fig. 3a). Differential abundance analysis (Fig. 3b) and composition bar plots (Fig. 3b) showed that term birth VMBs in older B/AA women had more pronounced enrichment of health-associated *Lactobacillus* species, e.g., *L. gasseri* and *L. jensenii*, and a greater reduction in dysbiosis-associated taxa, e.g., *G. vaginalis* and *P. amnii*. It was unexpected to find a lower abundance of protective *Lactobacillus* in term birth women among B/AA individuals aged 17-25 (Fig. 3b). However, this phenomenon was observed exclusively in the PRJNA393472 cohort (Fig. S7a-e) and not in the PRJNA725416 cohort (Fig. S7f-i).

**Fig. 3.**
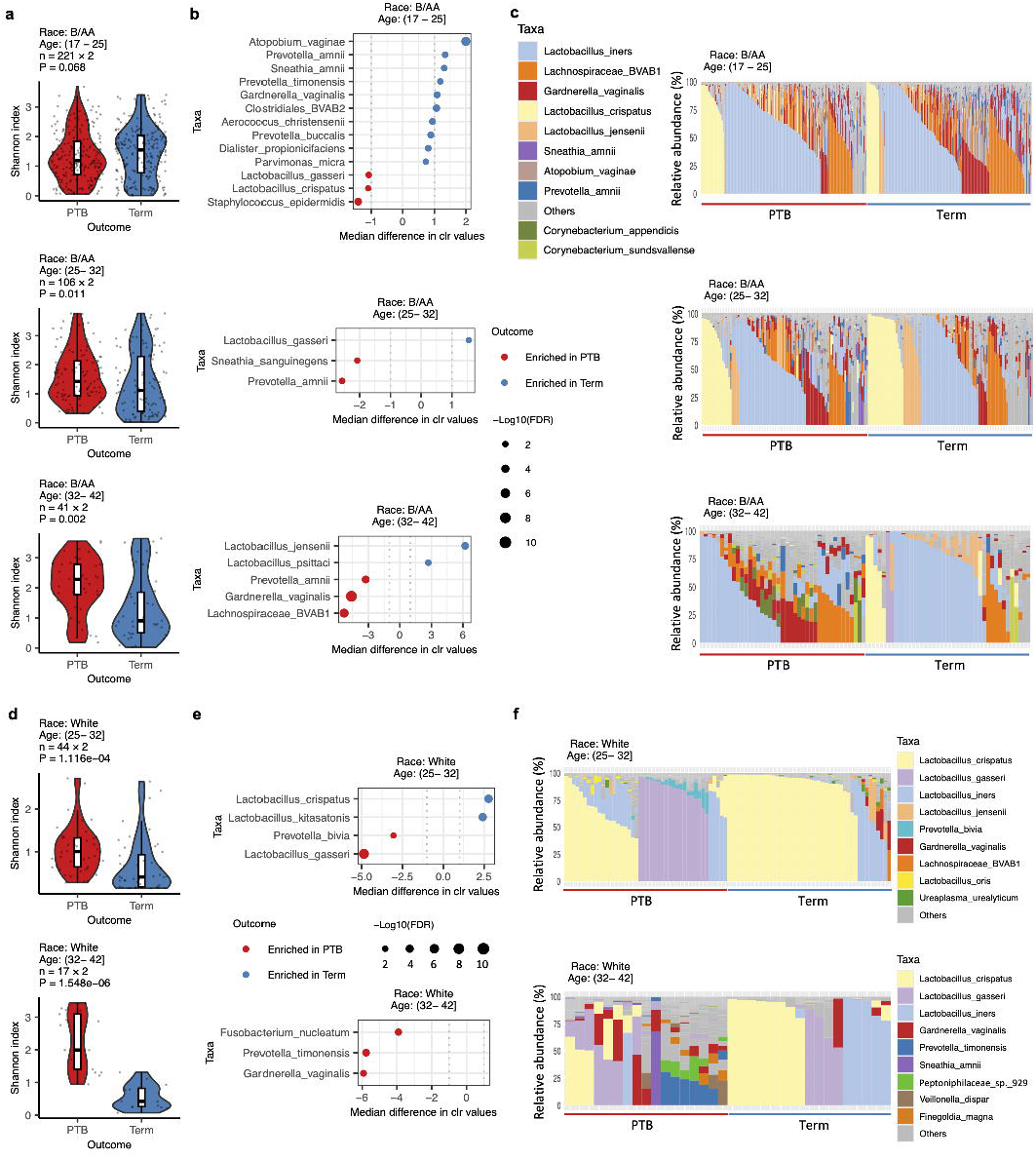
The impact of age on the association between the VMB and PTB in different racial groups. The VMB samples were stratified by race and age. Within each subgroup, PTB and term birth samples were case-matched based on BioProject, age, and gestational age (see Fig. S6). Differences in alpha diversity **(a)**, differential taxa abundances **(b)**, and VMB composition **(c)** between PTB and term birth in B/AA women across different age groups are shown. Differences in alpha diversity **(d)**, differential taxa abundances **(e)**, and VMB composition **(f)** between PTB and term birth in White women across different age groups are depicted. The results for Asian women aged 32.1-42 are shown in Fig. S9. Subgroup analyses for other White and Asian women are excluded due to fewer than 10 sample pairs. Statistical analyses included the two-sided Mann-Whitney U test for alpha diversity, the Adonis test for beta diversity, and the ALDEx2 package in R for differential relative abundance analysis. Lines in the boxplots represent maximum, 75% quantile, median, 25 quantile, and minimum values from top to bottom.

Similarly, term birth White women exhibited significantly lower alpha diversity in their VMBs, with this effect more pronounced in older age groups (Fig. 3d). Notably, VMBs in the 25.1-32 age group had higher levels of *L. crispatus* and lower amounts of *L. gasseri* (Fig. 3e and 3f). Further analysis showed that VMBs dominated by over 50% *L. crispatus* had lower levels of alpha diversity compared to those dominated by over 50% *L. gasseri* or *L. jensenii* (Fig. S8). This finding partially explains the reduced alpha diversity observed in term birth VMBs of White women aged 25.1-32 and implies that *L. crispatus* may be more effective than *L. gasseri* or *L. jensenii* in maintaining VMB homeostasis.

Term birth Asian women aged 32.1-42 also showed significantly lower alpha diversity, along with enrichment of health-associated *Lactobacillus* species and a reduction in dysbiosis-associated taxa (Fig. S9). Due to the limited number of cases (fewer than 10 pairs), results of other age groups are not shown.

### Interaction between age and preterm birth in relation to vaginal bacterial taxa

Given the evident role of age in mediating the association between the VMB and PTB in all the studied races, mixed-effects models were constructed to evaluate how the interaction between age and PTB affects the relative abundance of VMB taxa using the case-matched dataset as shown in Fig. S4d. The relative abundances were quantified using CLR normalized values. Results for the most abundant vaginal taxa are shown in Fig. 4, with the complete dataset available in Supplementary Dataset 1.

**Fig. 4.**
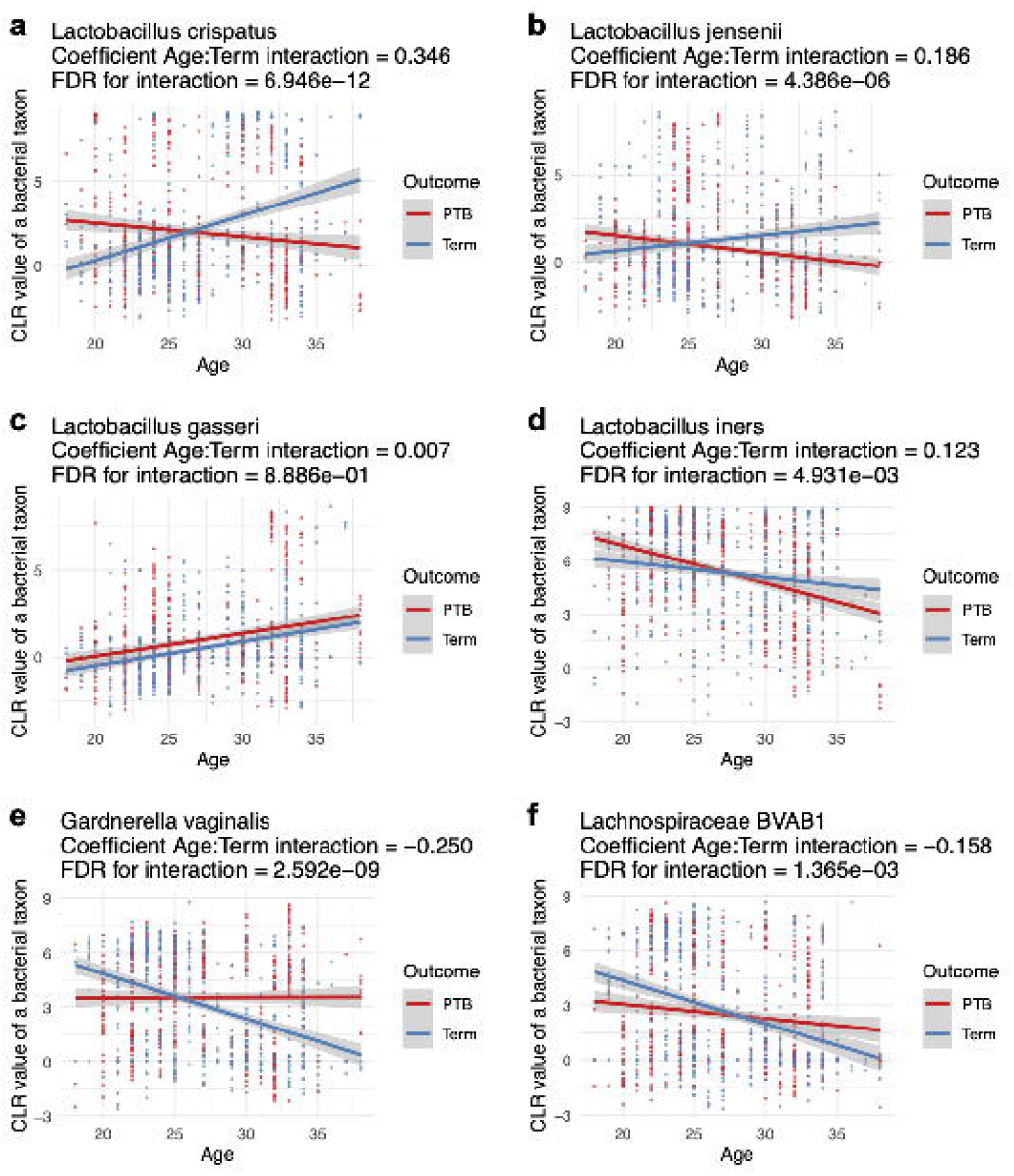
Interaction between age and preterm birth in relation to vaginal bacterial taxa. A mixed-effects model was implemented using the lm function from the lme4 package to test the interaction between age and pregnancy outcome on a specific vaginal taxon. The coefficients representing the interaction, along with the false discovery rates (FDRs) of the interaction, for *L. crispatus* **(a)**, *L. jensenii* **(b)**, *L. gasseri* **(c)**, *L. iners* **(d)**, *G. vaginalis* **(e)**, and *Lachnospiraceae* BVAB1 **(f)** are shown. A comprehensive list can be found in Supplementary Dataset 1.

The relative abundances of *L. crispatus* (Fig. 4a) and *L. jensenii* (Fig. 4b) increased with age in term birth women but decreased in PTB women. Positive interaction coefficients between age and delivery outcome indicate that for each 1-unit increase in age, the relative abundances of *L. crispatus* and *L. jensenii* increased by 0.346 and 0.186 units more, respectively, in term births compared to PTBs. In contrast, the relative abundance of *L. gasseri* increased with age in both term and PTB women, with no significant difference in the rate of increase (Fig. 4c). For *L. iners, G. vaginalis*, and *Lachnospiraceae* BVAB1, abundance decreased with age (Fig. 4d-f). However, the decline in *L. iners* was slower in term births, while decreases in *G. vaginalis* and *Lachnospiraceae* BVAB1 were more pronounced in term births compared to PTBs. The results for *L. crispatus, L. jensenii, G. vaginalis*, and *Lachnospiraceae* BVAB1 suggest that the overall status of the VMB tends to be more optimal with age in term births but declines with age in PTB.

### Association between the vaginal microbiota and preterm mediated by gestational age

The dbRDA analysis revealed that race, age, and gestational age had a greater impact on the VMB than birth outcomes (Fig. 1c). The influence of race and age on the relationship between PTB and the VMB has also been investigated. Following this approach, VMBs were further stratified by gestational age and case-matched for PTB and term birth based on BioProject, race, age, and gestational age (Fig. S10).

Previous studies reported significantly lower alpha diversity in the VMB during early pregnancy, particularly in the first trimester, for term births compared to PTB^18–20^. Our analysis was consistent with these findings, showing reduced alpha diversity in term births compared to PTB, with a more pronounced difference in early pregnancy (P-values for gestational age 44-135 days vs. later gestational ages in both B/AA (Fig. 5a) and White (Fig. 5b) women). Unlike earlier studies that found no significant lower levels of alpha diversity in term births in late pregnancy^19,20^, our data indicated a lower alpha diversity in late pregnancy for term births, but only among older B/AA women (≥25.1 years, Fig. 5a). Due to limited cases among White and Asian participants, comparisons were restricted to White women aged 25.1-32. Since there were more pronounced differences between PTB and term births in White compared to B/AA women (Fig. 2), it was not surprising that term births also had lower levels of alpha diversity than PTB in White women aged 25.1-32.

**Fig. 5.**
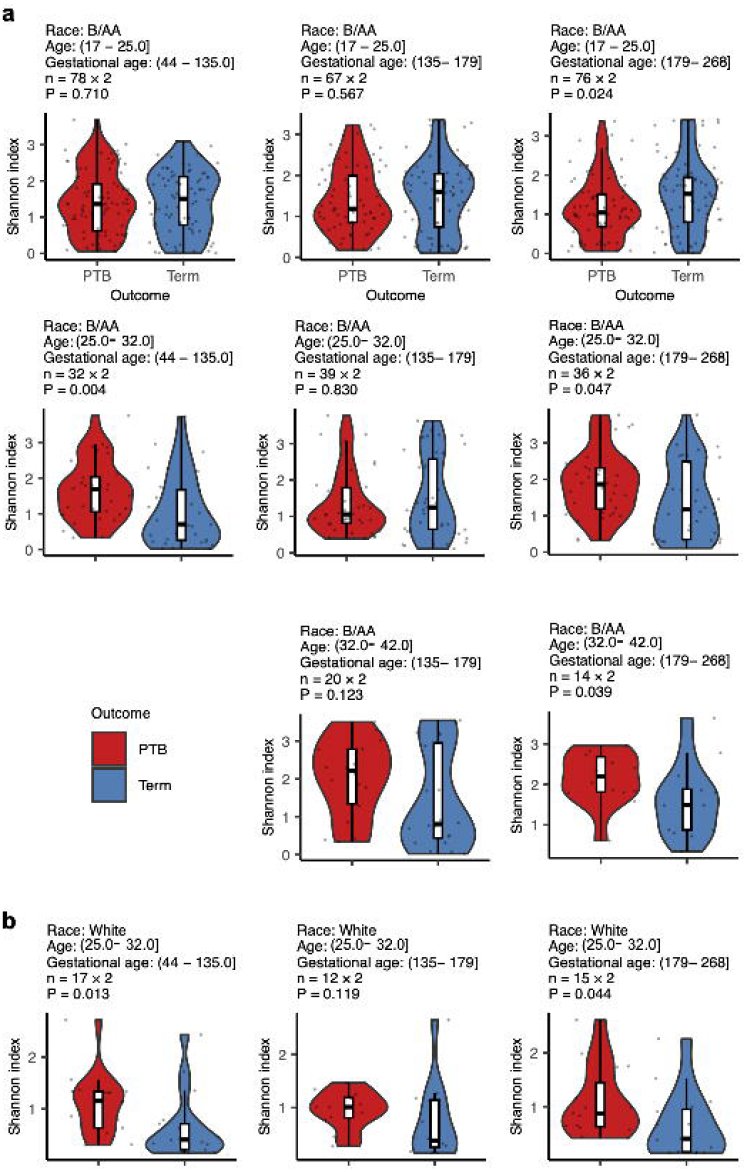
The impact of gestational age on the association between the VMB and PTB in different racial and age groups. The VMB samples were stratified by race, age, and gestational age. Within each subgroup, PTB and term birth samples were case-matched based on BioProject, age, and gestational age (see Fig. S10). Differences in alpha diversity between PTB and term birth in B/AA **(a)** and White **(b)** women across different age and gestational age groups are shown. The two-sided Mann-Whitney U test was applied for the comparison between PTB and term birth. Subgroups with sample pairs fewer than 10 are not shown. Lines in the boxplots represent maximum, 75% quantile, median, 25 quantile, and minimum values from top to bottom.

## Discussion

This study highlights the novel role of maternal age in mediating the association between the VMB and PTB. Age was identified as the second most influential factor affecting VMB composition, after race. Consistent with this finding, a previous study^17^ identified age as the fourth most important factor influencing the VMB, following poverty level, education, and marital status, all of which are linked to race^10^. Our data also showed that older women had more optimal VMB profiles, enriched in health-associated taxa, consistent with findings linking maternal age to a more optimal VMB^21^. Interestingly, the interaction between age and PTB revealed that while the relative abundance of *L. crispatus* and *L. jensenii* increased with age in term births, it decreased in PTBs, indicating a more protective role of these vaginal taxa in older term birth women in reproductive ages. Additionally, lower alpha diversity in the VMB of term pregnancies in late pregnancy was observed only in women aged 25.1 years or older. These findings highlight the complexity of age-related VMB changes and suggest that maternal age should be considered a critical factor in future studies on PTB and the VMB.

Previous studies have shown that estrogen is essential for increasing the abundance of protective *Lactobacillus* and maintaining overall vaginal health^1^. However, studies have not found higher estrogen levels in women aged 25.1 years or older compared to those under 25^22,23^. Therefore, estrogen levels cannot explain the relationship between age and changes in the abundance of vaginal taxa. Additionally, there is currently no evidence to suggest that younger women experience more severe fluctuations in the composition of the VMB due to hormonal changes during reproductive years or that they possess a less effective immune system in maintaining VMB homeostasis. A plausible hypothesis is that age-related changes in behavioral and lifestyle factors may influence the VMB. For instance, more frequent sexual activity among younger women^24^ could modulate the VMB, potentially resulting in a less stable or optimal VMB. However, the more pronounced differences in the VMB between PTB and term birth observed in older reproductive-age women could be attributed to a less efficient immune system in this age group. As immune system function declines with age^25^, maintaining an optimal VMB may become more critical in reducing the risk of PTB in older women of reproductive age.

The difference in the VMB between PTB and term birth becomes notably less significant when cases are matched based on BioProject, race, age, and gestational age, compared to analyses without a case-matching design. Combined with the results of the dbRDA analysis, this highlights the critical role of race, age, and gestational age in shaping the VMB. These factors should therefore be carefully accounted for in studies investigating associations between the VMB and PTB.

It is well established that an optimal VMB dominated by *Lactobacillus* species is associated with a lower risk of PTB. However, our data revealed a lower abundance of protective *Lactobacillus* in term-birth women within one specific group, B/AA aged 17-25. Notably, this observation was inconsistent across the two cohorts studied, PRJNA393472 and PRJNA725416, suggesting that additional factors may have influenced the results observed in the PRJNA393472 cohort for this subgroup. Due to limitations in the metadata available across all studied cohorts, this analysis included only three maternal characteristics: race, age, and gestational age. Incorporating more comprehensive metadata in future studies could offer a deeper understanding of the role of the VMB in PTB.

A significant limitation of this study is the disparity in case numbers across racial groups, which undermines the statistical power for certain subgroups. For example, Asian women across all ages and White women under 25 years had limited sample sizes. Although similar trends of differences between PTB and term birth were observed in these subgroups (data not shown), the robustness of statistical comparisons was compromised, making the results less reliable.

## Data availability

Publicly available datasets were downloaded from the NCBI database (https://www.ncbi.nlm.nih.gov/). The BioProject IDs with data used in this study are listed below: PRJNA300860, PRJNA393472, PRJNA687274, and PRJNA725416.

## Code availability

The underlying code for this study can be accessed via this link [https://github.com/xyun1275/VMB_PTB/blob/main/results_visualization.R].

## Ethics statement

Study protocols were approved by the Northwest Women’s and Children’s Hospital Institutional Review Board (IRB) under protocol number IRB 2024-100.

## Author contributions

Y.X.: Funding acquisition, Investigation, Methodology, Writing-review & editing. Q.W.: Methodology, Writing-original draft. D.L.: Data curation, Methodology. F.J.: Data curation, Methodology. F.C.: Project administration, Resources. Z.Z.: Investigation, Methodology. Y.S.: Data curation, Methodology. S.W.: Project administration, Supervision, Funding acquisition, Writing-review & editing. All authors read and approved the final manuscript.

## Funding

This work was supported by Innovation Capability Strong Foundation Plan of Xi’an City (Medical Research Project, 22YXYJ0117), Key Research and Development Program of Shaanxi (Program No. 2023-YBSF-320), 2024 Shaanxi Provincial Health high-level talent (team) cultivation plan-Young Talent Project, and the Fundamental Research Funds for the Central Universities (xzy012024146), National Natural Science Foundation of China (32170114 and 31770152), and Shaanxi Fundamental Science Research Project for Chemistry & Biology (Program No. 22JHZ008).

## Acknowledgments

We acknowledge all project investigators from University of Sheffield, Stanford University School of Medicine, Mayo Clinic, and Emory University for their contributions to generating the publicly shared data. We also thank the participants involved in sample collection for these BioProjects.

## Conflict of interest

All authors declare no financial or non-financial competing interests.

## Notes

### Competing Interest Statement

The authors have declared no competing interest.

