## Supplementary materials for "Role of Age in Mediating the Association Between the Vaginal Microbiota and Preterm Birth"

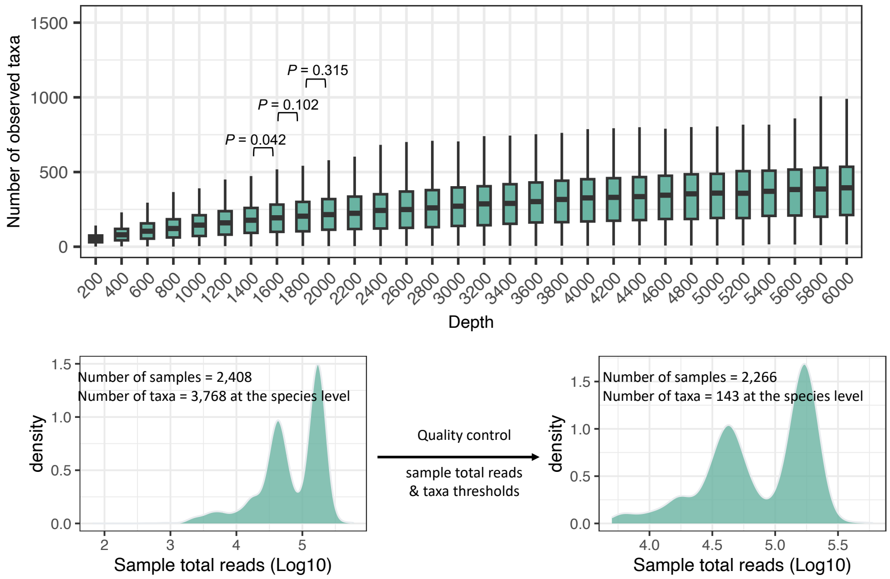


**Fig. S1 Quality control of the VMB. (a)** Alpha rarefaction analysis to determine the threshold for a sample's total reads. Difference between two rarefaction depth was quantified by the two-sided Mann-Whitney U test. **(b)** The numbers of samples and vaginal taxa included in the study, both before and after quality control, are shown. Quality control was performed based on sample total read counts and taxa thresholds as detailed in the Methods section.

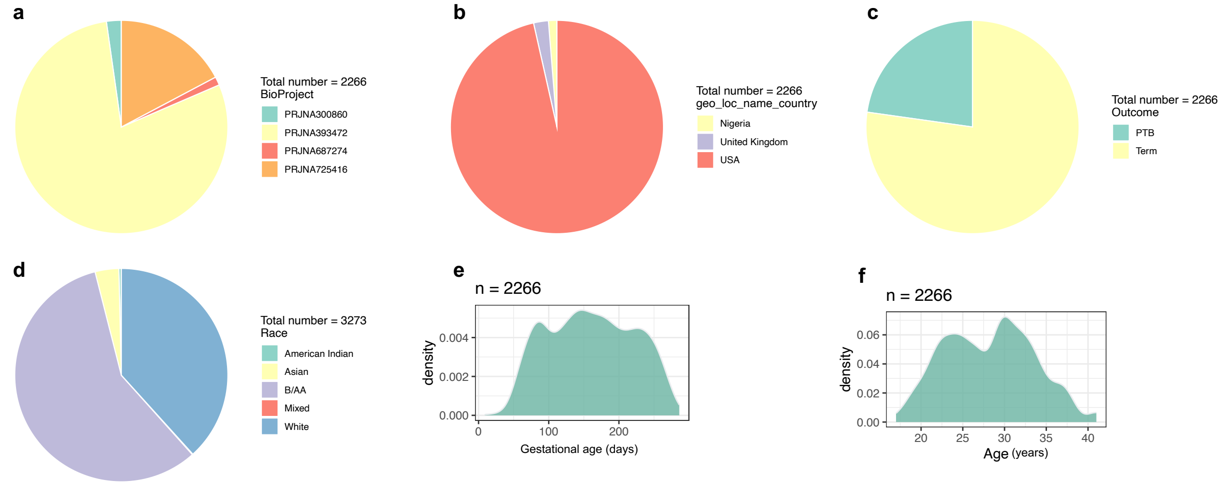


**Fig. S2 The metadata profiles of the VMBs.** Distribution of the VMBs in different BioProjects **(a)**, Countries **(b)**, pregnancy outcomes **(c)**, races **(d)**, gestational ages **(e)**, and maternal ages **(f)** are shown.

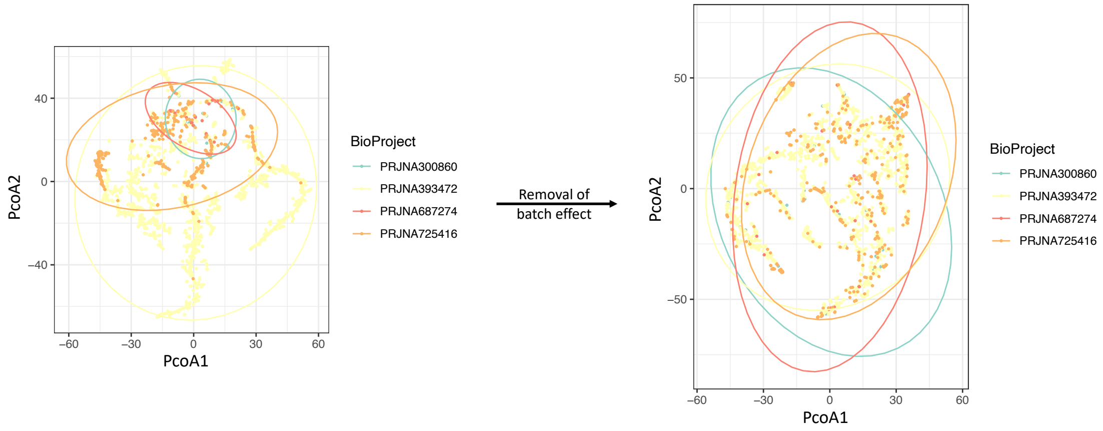


**Fig. S3 Beta diversity of the VMBs before and after batch-effect removal are visualized by PCoA plots.**

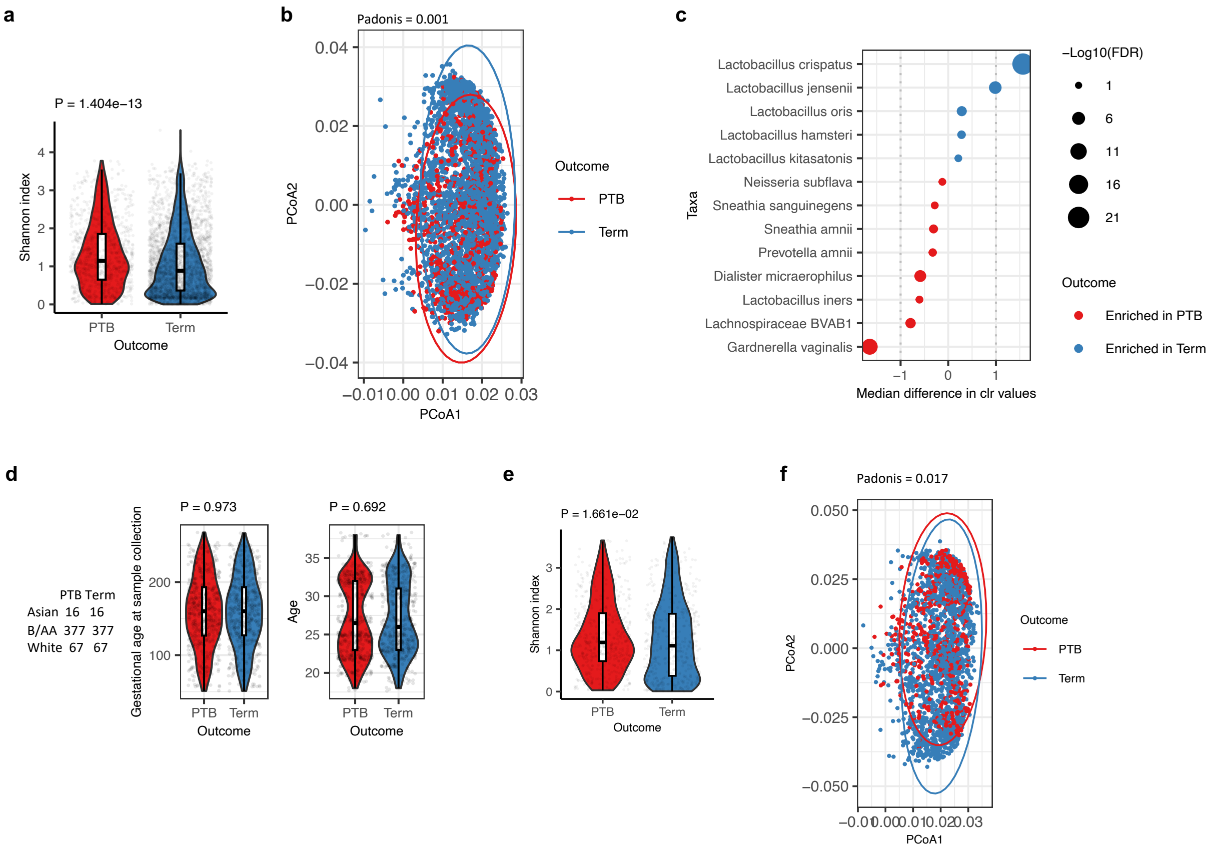


**Fig. S4 The association between the VMB and PTB in cohorts with and without a case-matched design.** Samples collected from the four BioProjects described in Fig. S2a were used to measure the differences between the VMBs of PTB and term birth in alpha diversity **(a)**, beta diversity **(b)**, and differential abundance changes **(c)**. The VMBs from PTB and term birth cases were matched based on race, BioProject, and similar age and gestational age **(d)**. The differences between the VMBs of PTB and term birth in alpha diversity **(e)** and beta diversity **(f)** are shown. Statistical analyses included the two-sided Mann-Whitney U test for alpha diversity, the Adonis test for beta diversity, and the ALDEx2 package in R for differential relative abundance analysis.

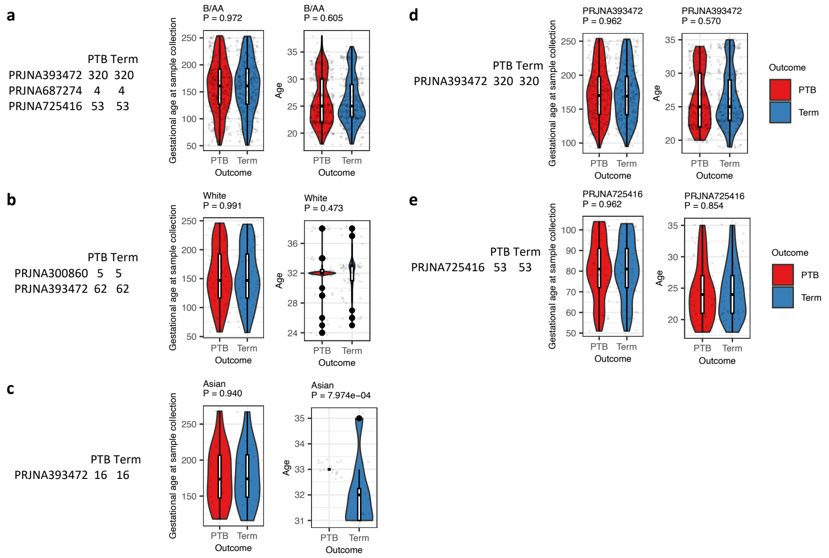


**Fig. S5 Case match design for investigating the impact of race on the association between the VMB and PTB.** The VMB samples were categorized into three racial subgroups, i.e., B/AA, White, and Asian. In each subgroup, PTB and term birth samples were case-matched by BioProject, age, and gestational age. Comparisons of gestational age and maternal age using the two-sided Mann-Whitney U test are presented for B/AA **(a)**, White **(b)**, and Asian **(c)** subgroups, respectively. Further analysis tested differences in alpha diversity between PTB and term birth in B/AA women across two individual cohorts. Case matching for PRJNA393472 **(d)** and PRJNA725416 **(e)**, based on BioProject, age, and gestational age, is shown.

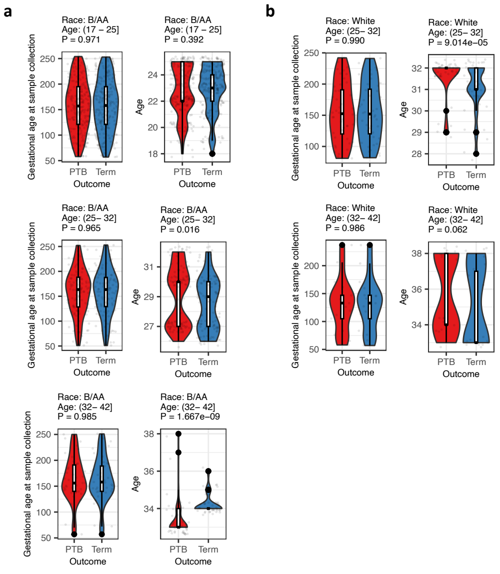


**Fig. S6 Case match design for investigating the impact of age on the association between the VMB and PTB in different racial groups.** The VMB samples were stratified by race and age. In each subgroup, PTB and term birth samples were case-matched by BioProject, age, and gestational age. Comparisons of gestational age and maternal age between PTB and term birth using the two-sided Mann-Whitney U test are presented for B/AA **(a)** and White **(b)** subgroups, respectively.

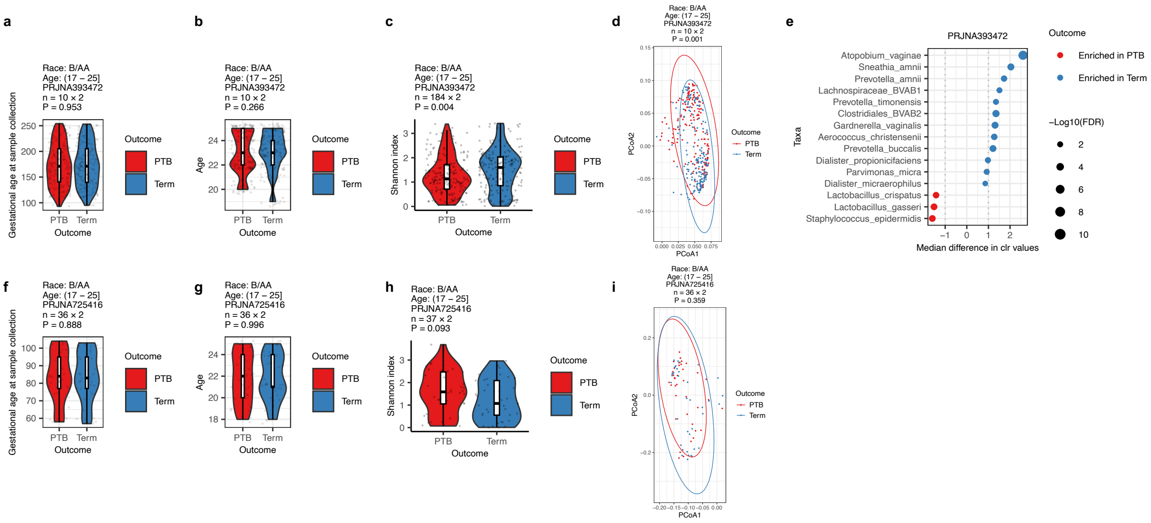


**Fig. S7 Difference between PTB and term birth in B/AA aged 17-25 in two individual cohorts.** Case match between PTB and term birth on gestational age **(a)** and age **(b)** for the PRJNA393472 cohort is shown. The differences between the VMBs of PTB and term birth in alpha diversity **(c)**, beta diversity **(d)**, and differential abundance changes **(d)** in the PRJNA393472 cohort are illustrated. Case match between PTB and term birth on gestational age **(f)** and age **(g)** for the PRJNA725416 cohort is shown. The differences between the VMBs of PTB and term birth in alpha diversity **(h)** and beta diversity **(i)** in the PRJNA725416 cohort are illustrated. Statistical analyses included the two-sided Mann-Whitney U test for alpha diversity, the Adonis test for beta diversity, and the ALDEx2 package in R for differential relative abundance analysis.

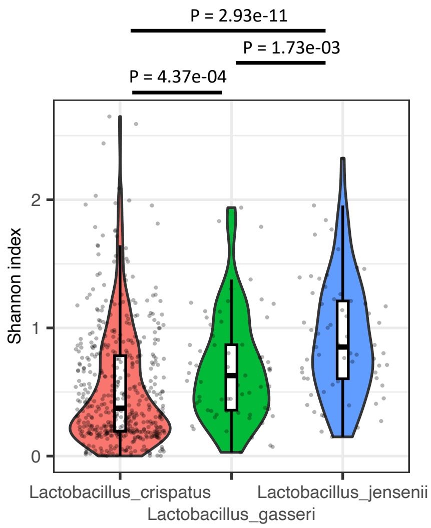


**Fig. S8 Alpha diversity of the VMB dominated by different *Lactobacillus* species.** The predominance of a specific *Lactobacillus* species in the VMB was defined as having a relative abundance exceeding 50%. Differences in alpha diversity between two VMB groups were assessed using the two-sided Mann-Whitney U test.

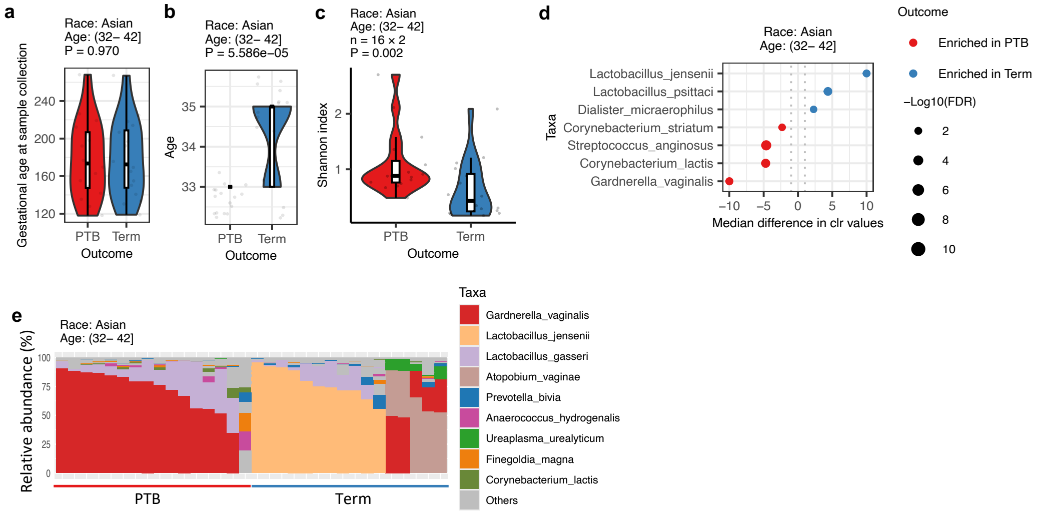


**Fig. S9 Difference between PTB and term birth in Asian aged 32.1-42.** Case match between PTB and term birth on gestational age **(a)** and age **(b)** is shown. The differences between the VMBs of PTB and term birth in alpha diversity **(c)**, differential abundance changes **(d)**, and the composition of the VMB are illustrated. Statistical analyses included the two-sided Mann-Whitney U test for alpha diversity, the Adonis test for beta diversity, and the ALDEx2 package in R for differential relative abundance analysis.

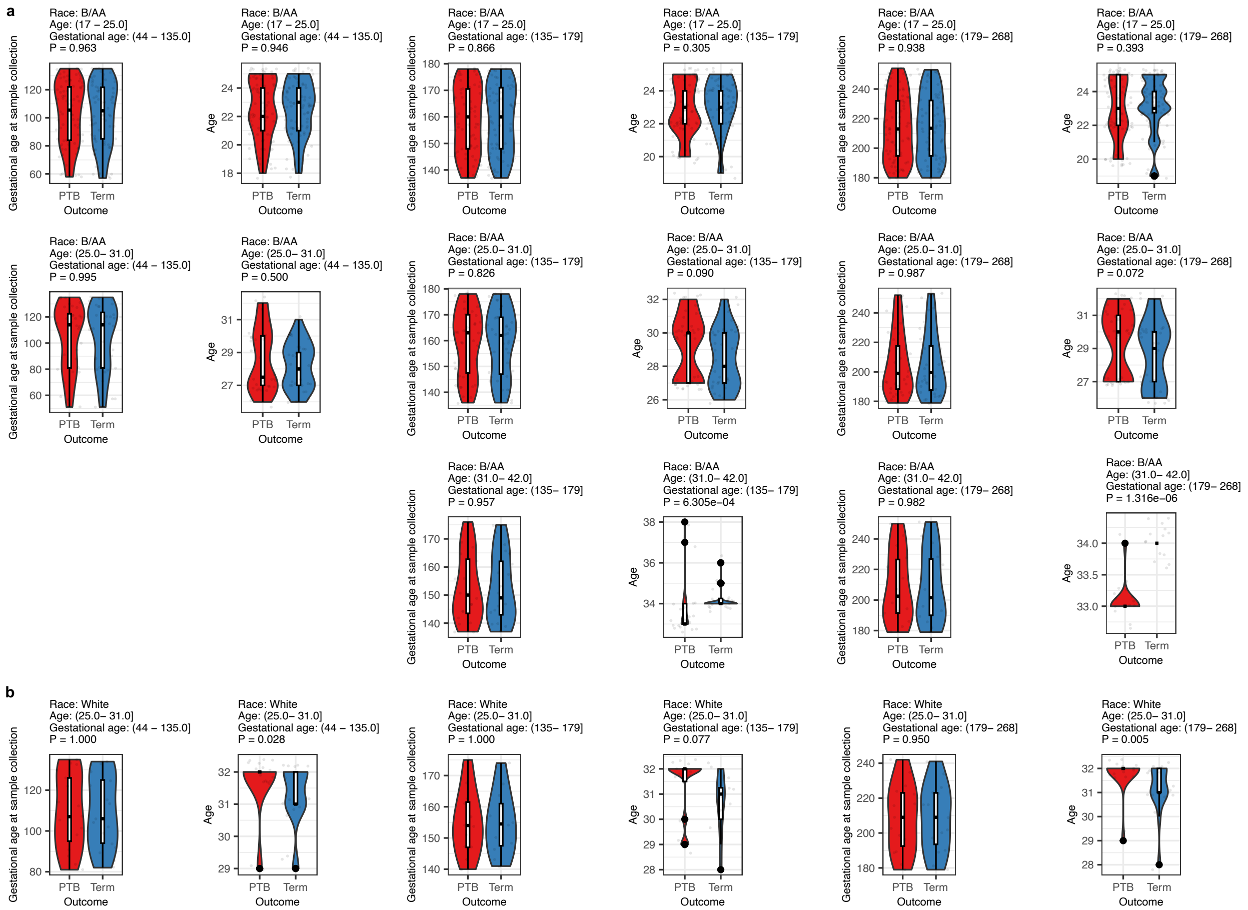

**Fig. S10 Case match design for investigating the impact of gestational age on the association between the VMB and PTB in different racial and age groups.** The VMB samples were stratified by race, age, and gestational age. In each subgroup, PTB and term birth samples were case-matched by BioProject, age, and gestational age. Comparisons of gestational age and maternal age between PTB and term birth using the two-sided Mann-Whitney U test are presented for B/AA **(a)** and White **(b)** subgroups, respectively.
